# *PySOFI*: an open source Python package for SOFI

**DOI:** 10.1101/2021.10.16.464651

**Authors:** Yuting Miao, Shimon Weiss, Xiyu Yi

## Abstract

Super-resolution optical fluctuation imaging (SOFI) is a highly democratizable technique that provides optical super-resolution (SR) without requirement of sophisticated imaging instruments. An open source package for SOFI algorithm is needed to support not only the utilization of SOFI, but also the community adoption and participation for further development of SOFI. In this work, we developed *PySOFI*, an open source python package for SOFI analysis that offers the flexibility to inspect, test, modify, improve and extend the algorithm. We provide a complete documentation for the package and a collection of Jupyter Notebooks to demonstrate the usage of the package. We discuss the architecture of *PySOFI*, illustrate how to use each functional module, and demonstrate how to extend the *PySOFI* package with additional modules. We expect *PySOFI* to facilitate efficient adoption, testing, modification, dissemination and prototyping of new SOFI-relevant algorithms.

## 1. INTRODUCTION

Super-resolution optical fluctuation imaging (SOFI) [1] is a widely used optical super-resolution method applicable for a broad range of conditions, where sophisticated control on the instrument and sample preparations are not required. It has attracted a growing community of active practitioners and developers over a decade. The advancements utilizing this technology include innovations in blinking dyes and fluorescent proteins, sample preparation [2, 3, 4, 5, 6], illumination schemes, experiment designs, data processing methods [7, 8, 9, 10, 11, 12, 13, 14, 15], and integration with other methods [16, 17, 18, 19, 20, 21]. With *n*^*th*^ order cumulant analysis, the theoretical resolution improvement of SOFI is 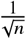. This limit increases to 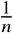 when combined with deconvolution, presenting a great potential for further advancements for SOFI [1].

Our previous work have identified imperfections in high-order SOFI cumulants (e.g., cusp-artifacts [22]), where we introduced the concept of *virtual emitters* to interpret the physical meaning of SOFI cumulants, and identified the origin of such artifacts to be the adjacent negative and positive *virtual emitters* presented in the high-order SOFI cumulant image [22]. We also demonstrated how the validity of one of the most widely used SOFI processing method, bSOFI is negatively impacted [7].

We believe an insightful and thorough understanding of the method is crucial to ensure solid advancements in both SOFI and SOFI-relevant innovations. However, for new investigators without prior experience with SOFI analysis, there is often a steep learning curve to fully understand, modify, and extend the existing open-source packages [7, 23]. The existing SOFI analysis routines are implemented in either ImageJ or MATLAB. ImageJ requires professional programming skills if customization and modifications are required, while MATLAB requires a paid license. Such limitations present a greater challenge for new investigators who are interested in joining the SOFI community but prefer not to use the existing packages blindly.

Here we present *PySOFI*, an open source package for SOFI analysis implemented in Python. Benefited from the active open-source community and the abundance of free learning materials for Python, *PySOFI* offers an easy option for investigators interested in adopting the SOFI algorithm. *PySOFI* focuses on engaging the community and is designed to be simple, modular, and highly customizable. *PySOFI* is hosted on GitHub to facilitate utilization, improvements, and continuous maintenance by the interested users and developers. A collection of examples is provided in the form of Jupyter Notebooks. One can use *PySOFI* to explore and characterize SOFI analysis, validate the results from the prior studies, and gain insights through exploration. *PySOFI* is also useful for the prototyping of new methods to extend SOFI algorithm. Similar Jupyter Notebooks can be adapted to promote the new methods and improve the reproducibility of the results. We expect *PySOFI* to appeal to both beginners and experts, it facilitates innovations where modification and extensions are required, and further promote the scientific advancements among scientists interested in SOFI.

The rest of the manuscript is organized as follows. Section 2 provides an overview of the *PySOFI* package. Section 3 discusses the *PySOFI* software architecture design and analysis pipeline, together with analysis examples for various of modules. Section 4 provides an example of extending *PySOFI* with an additional analysis module called *MOCA*. Section 5 summarizes the work and discusses future directions.

## 2. *PYSOFI* OVERVIEW

We designed a straightforward architecture for the *PySOFI* package. As shown in (figure 1), *PySOFI* contains eight independent function modules (in the `functions` folder) and one data class (`PysofiData`). A detailed description of *PySOFI* is available in our [online documentation](https://xiyuyi-at-llnl.github.io/pysofi/build/html/). To get started with the installation, the user can follow [this page](https://xiyuyi-at-llnl.github.io/pysofi/build/html/start.html).

**Figure 1:**
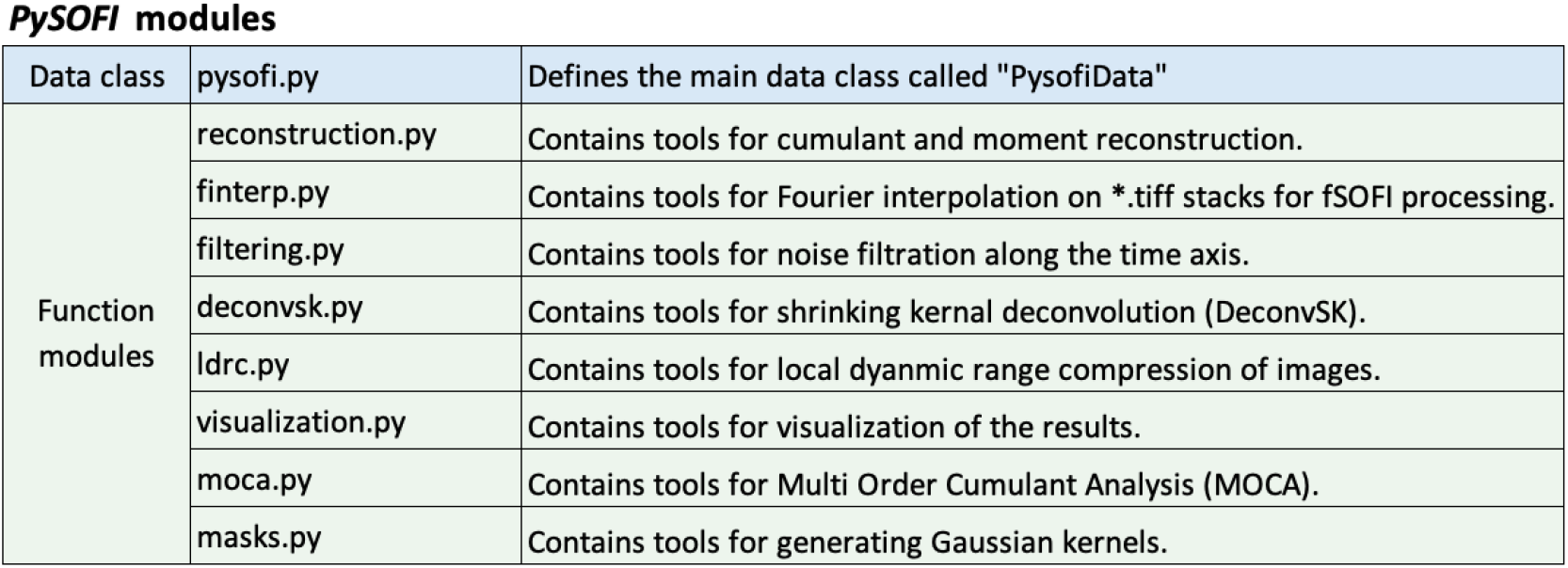
*PySOFI* modules. *PySOFI* contains one data class and eight function modules. Detailed descriptions are available in the [online documentation](https://xiyuyi-at-llnl.github.io/pysofi/build/html/) for *PySOFI*.

The following modules are implemented to facilitate the *PySOFI* analysis pipeline. The `reconstruction.py` module provides capabilities for SOFI moments and cumulants calculations [1], as well as bleaching correction for a TIFF movie. The `finterp.py` module provides Fourier interpolation on a TIFF stack, which is a necessary step for fSOFI alike analysis [9]. `filtering.py`, `deconvsk.py`, and `ldrc.py` constitute a collection of modules relevant to SOFI 2.0 [11] analysis. Specifically, the `filtering.py` module is for pixel-wise noise filtering along the time axis, the `deconvsk.py` module is for shrinking kernel deconvolution (DeconvSK)[10], and the `ldrc.py` is for local dynamic range compression (*ldrc*) of images with a large dynamic range of pixel values [10]. Additionally, the `moca.py` module provides capabilities for multi-order cumulant analysis (MOCA). The `masks.py` module is used to generate Gaussian kernels, and the `visualization.py` module provides visualization options using an interactive visualization package `bokeh`. The data class module (`PysofiData`) is encapsulated in the `pysofi.py` file. The input parameters from the users, the raw data, and the intermediate results are bundled in the `PysofiData` object as attributes, and the processing steps as methods. The processing steps are implemented as function modules, and imported and used in the data class module. In summary, the specific functions are implemented in the function modules, while `PysofiData` serves the purpose of organizing the data processing workflow.

Figure (2) provides the data-flow diagram that demonstrate the connections (arrows) between different processing steps (green squares) and different types of data (purple ovals). Three collections of SOFI analysis routines are implemented in the `PysofiData` class, including the “Shared Processes” that contains the traditional SOFI analysis steps [1], the “SOFI 2.0” collection that contains the routines for SOFI 2.0 processing [11] and the “MOCA” collection that enables multi-order cumulant analysis (MOCA) [10]. In the “Shared Processes” block, the processing steps including bleaching correction (BC), Fourier interpolation (FI), and moment and cumulant calculations. The processing steps can be performed in various sequences (green arrows). In the “SOFI 2.0” block, one can perform noise filtering, deconvolution (DeconvSK), and local dynamic range compression (ldrc) on the image. The “MOCA” collection can be used to estimate the blinking statistics distribution (*ρ*-map) and brightness distribution (*ϵ*-map). In the data processing workflow, one can save and load the intermediate results for each processing step (purple arrows). For example, in the “Shared Processes” collection, the intermediate results (purple ovals) can be saved as separate new TIFF files or stored as attributes in the `PysofiData` class object, and then passed to another processing step (purple arrow).

**Figure 2:**
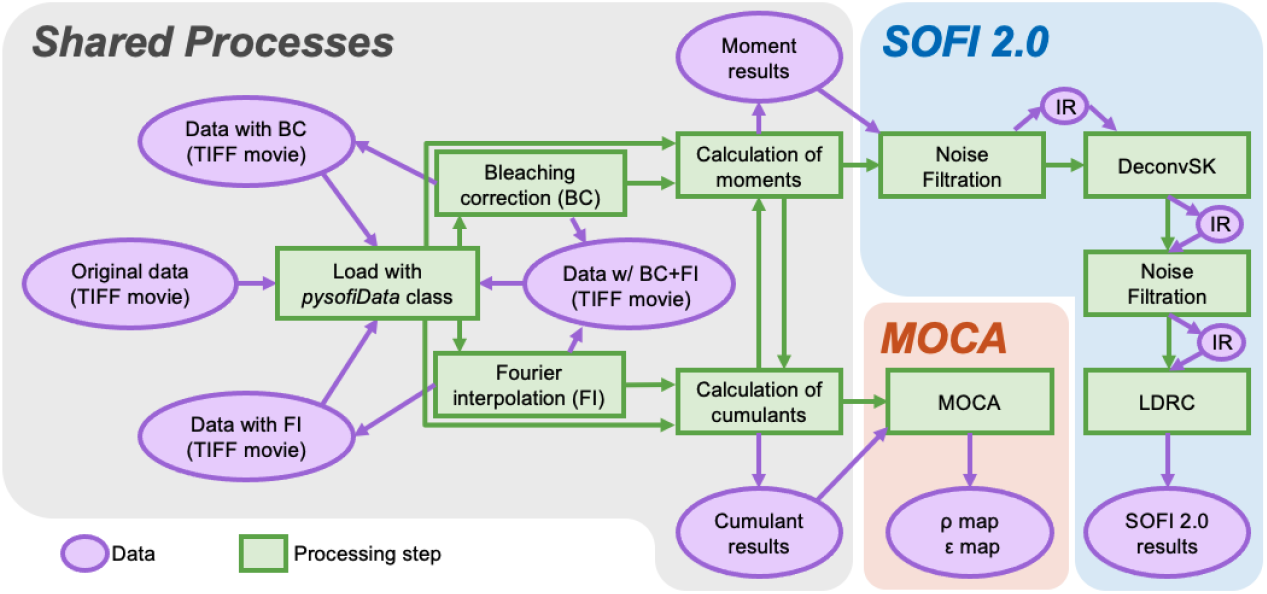
Data-flow diagram for *PySOFI*. Three collections of SOFI analysis routines are implemented in *PySOFI* as depicted in the diagram: Shared Processes, SOFI 2.0 analysis, and MOCA analysis. Green squares represent data processing steps with functionalities labeled for each step. The purple ovals represent the data types as labeled in the diagram. Intermediate results are abbreviated as “IR”. The green arrow represents the direction of the data flow between different steps, and the purple oval represents input and output data types at different processing steps.

In general, we adopted a simple architecture for *PySOFI* with a collection of independent function modules and only one class module (the data class). The functions are imported and used inside the data class across different methods as needed, therefore the implementation is flexible with minimum repetition of codes. The function modules can be implemented, modified, and tested independently, ensuring flexibility and convenience for maintenance. Extending the package can be done by implementing additional function modules. It can be used as a standalone process, or be integrated into the data processing workflow through the `PysofiData` class. Note that the “MOCA” collection is an implementation example that extends *PySOFI* package in a modular fashion. The investigators also have the flexibility to disseminate the *PySOFI* package and construct their own data processing workflow (similar to the `PysofiData` class).

## 3. IMPLEMENTATION OF SOFI ANALYSIS USING *PYSOFI*

We provide a collection of Jupyter Notebooks (outlined in Figure 3) as examples for *PySOFI* implementations and applications. The prefix (E#) of each filename is used as a reference to each notebook in the following text for simplicity. We present example *PySOFI* analysis steps (E1 to E7), visualization of the result with combined color-map and transparency-map (E8), and the effect of data acquisition length on SOFI reconstruction performance (E9, E10). We also demonstrate SOFI 2.0 analysis (E11) and characterization of cusp-artifacts (E12 and E13). The analysis processes are integrated through the `PysofiData` class for all the notebooks except for the demonstration of noise filtration (E2).

**Figure 3:**
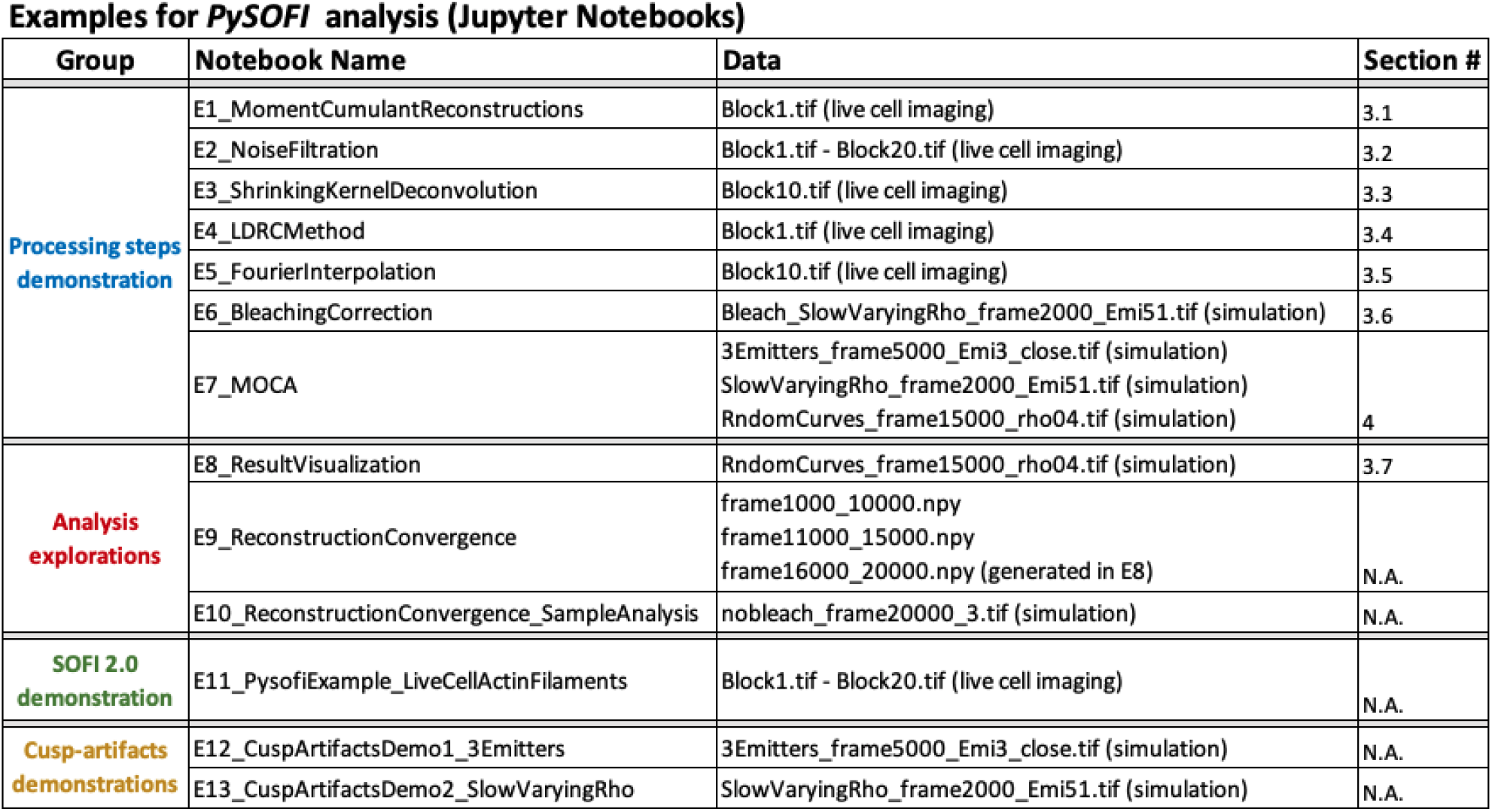
Jupyter notebook examples for *PySOFI*. We provide 13 *PySOFI* demonstrations as Jupyter Notebooks which can be categorized into to 4 Groups (first column). The filenames (second column) indicates the focus of each Jupyter Notebook. The relevant data sets (third column) are shared on [figshare](https://figshare.com/s/47d97a2df930380c96bb). Brief descriptions of some notebooks (E1 to E8) are provided in the relevant section (fourth column). The theory behind E9 to E13 are not included in this manuscript but the relevant concepts are discussed in [22] and [11]. The notebooks are the *PySOFI* implementations of the relevant methods to support the utilization of them. In particular, in E11, we show the general guidelines for performing SOFI 2.0 analysis on live-cell fluorescence imaging results using *PySOFI*.

In the text below, we provide brief descriptions of E1 to E6. The detailed description and examples are provided in the Jupyter Notebooks in the online [Github repository](https://xiyuyi-at-llnl.github.io/pysofi/build/html/demos.html).

### 3.1 Moment and cumulant reconstructions (E1)

Traditionally, SOFI achieves resolution enhancement by computing different orders of cumulants of optical signal fluctuations in time. The theoretical resolution enhancement for SOFI is 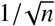 fold for the *n*^*th*^ order SOFI cumulant. Once combined with deconvolution, the theoretical resolution enhancement can increase to 1*/n*.

To obtain the *n*^*th*^ order SOFI cumulant, one way is to construct the *n*^*th*^ order cumulant as a polynomial consists of moments from the first order to the *n*^*th*^ order, as shown in the previous work [1]. Another way, which is used by *PySOFI*, is to construct the following recursive relation: 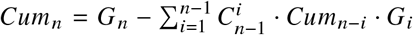, where *Cum*_*n*_ represents the *n*^*th*^ order cumulant, *G*_*n*_ represents the *n*^*th*^ order moment, and 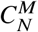 means the number of combinations of “N choose M”. Regarding the moment calculations, *PySOFI* support calculations of moments directly from the time series of each pixel. The moments can be also calculated as a reconstruction from a series of cumulants as used in our previous study [11].

The calculation of cumulants and moments are the fundamental processing elements in the SOFI analysis. The `PysofiData` class organizes the analysis workflow and can be used to calculate both moments and cumulants. Essentially, the relevant function modules are imported and integrated in the `PysofiData` to support such analysis. For example, the following scripts would calculate the 4^*th*^ order moment and cumulant of the specified TIFF stack named `Block1.tif` through the `PysofiData` class:

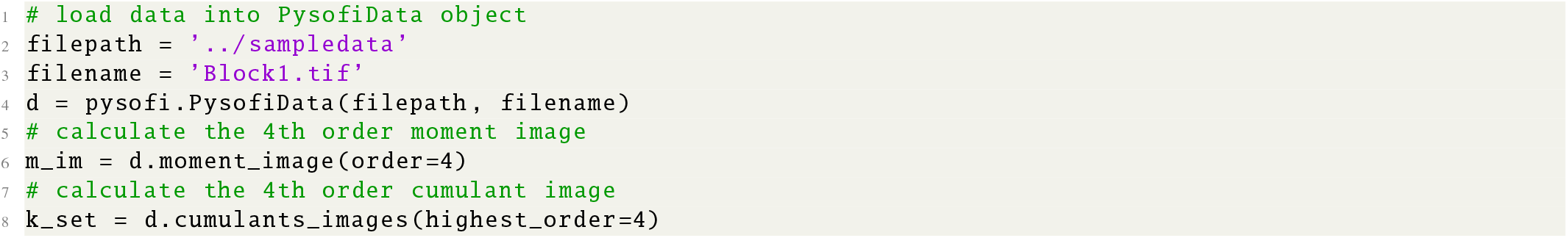

We can also directly import the function module, `reconstruction.py`, to perform the relevant calculations. This option is designed to support dissemination of the *PySOFI* package to facilitate independent analysis, which is often useful when developing new methods built upon SOFI analysis. The following scripts demonstrate how to perform such analysis with moment and cumulant calculations up to 4^*th*^ order:

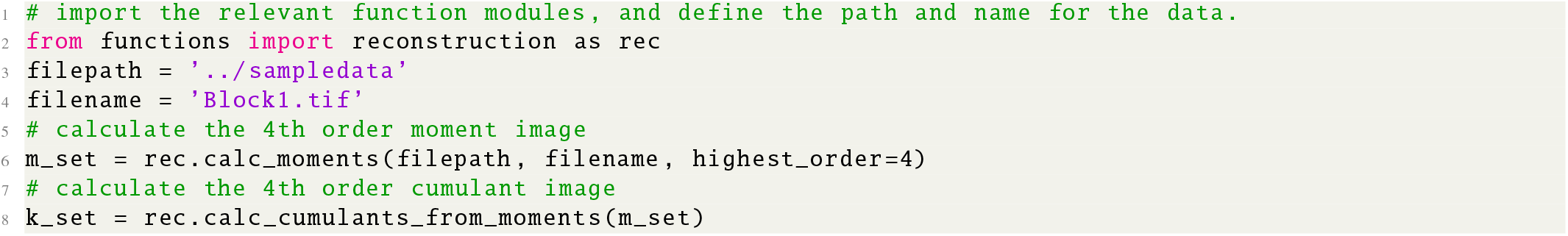

More detailed demonstrations are available in the corresponding Jupyter Notebook ([E1](https://github.com/xiyuyi-at-LLNL/pysofi/blob/master/Notebooks/E1_MomentCumulantReconstructions.ipynb)).

### 3.2 Temporal Noise Filtering (E2)

Temporal noise filtering is fundamental in the image processing for fluorescence microscopy, especially in scenarios where continuous and prolonged live cell imaging is desired where the excitation power is maintained at a low level to minimize photo toxicity and photo-bleaching. The lower excitation power often results in reduced signal to noise ratio. Traditional noise filtering is performed with a spatial filter where each image for every given time instance is spatially filtered independently. However, because noise filtering in the spatial spectrum domain is equivalent to a convolution operation of the image with the kernel corresponds to the inverse Fourier transform of the low-pass filter, it is conceivable that the spatial noise filtering would reduce the spatial resolution. On the other hand, to achieve a super-resolution movie, we are focusing on the sample conditions where the semi-static assumption is valid, which requires slow dynamics in the sample and the temporal noise filtering has been proven useful [11]. This is because slow dynamics ensures that the signal of interest exists in the low frequency domain while the noise is populated in the high frequency domain in the time axis, therefore the temporal spectrum filtering can be effective. Additionally, because this filtering is performed along the time axis, the spatial resolution is not directly influenced.

We have implemented such temporal noise filtering in *PySOFI* as a function module `filtering.py`. It is useful when analyzing multiple TIFF stacks corresponding to consecutive time-blocks. In such scenario, the feature is assumed to be semi-static within each individual time block, and the corresponding TIFF stack is analyzed independently. We can perform the temporal noise filtering on the results across all the time blocks to further enhance the image quality.

For example, we can perform the temporal noise filtering on the 6^*th*^ order moment images calculated from 20 blocks of TIFF stacks (each contains 200 frames) using the following scripts:

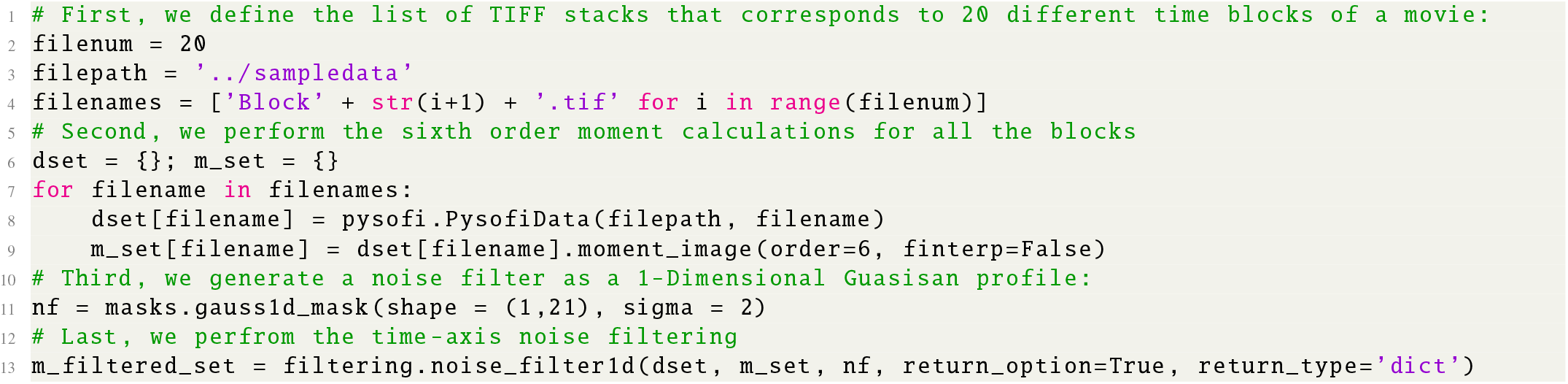

The results from the temporal noise filtering are stored as a dictionary in the `m_filtered_set`, where keys for elements are file names for each block of TIFF images, and values are the corresponding filtered images. The filtered images are also updated to each `PysofiData` objects as a `PysofiData.filtered` attribute. More detailed demonstrations are available in the corresponding Jupyter Notebook ([E2](https://github.com/xiyuyi-at-LLNL/pysofi/blob/master/Notebooks/E2_NoiseFiltration.ipynb)).

### 3.3 Shrinking kernel deconvolution - DeconvSK (E3)

With the help of high-order SOFI analysis, the point spread function (PSF) of the optical system can be estimated [8], and deconvolution can be used to further enhance the image resolution by a factor of 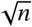 for *n*^*th*^ order SOFI cumulant [8]. In SOFI 2.0 [11, 10], we developed and applied the shrinking kernel deconvolution (DeconvSK) method to SOFI images. The principle of deconvSK is that for a system with a Gaussian PSF, the estimated convolution kernel can be decomposed into a series of smaller Gaussian kernels. We design the series in a way to include only one parameter, and form a series of Gaussian kernels with decreasing width. More specifically, denote the system PSF (2D Gaussian function) as *U*, we have

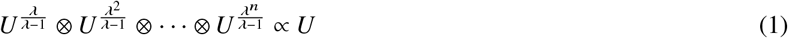

where *λ* is an empirical parameter between 1 and 2 that determines the exponent, ⊗ represents the convolution operation, and ∝ represents the proportionality. In this way, we can decompose the overall deconvolution task into a series of lighter deconvolution tasks.

In *PySOFI*, DeconvSK is implemented in the function module *deconvsk*.*py* as well as in the `PysofiData.deconvsk()` method. `DeconvSK` requires two different input parameters, the *λ* in equation 1 can be set to be between 1 to 2. The width of the overall PSF with Gaussian approximation can be obtained from the cross-cumulant analysis [8] or directly estimated from the instrument configurations. The DeconvSK processing can be performed with the following commands through a `PysofiData` object:

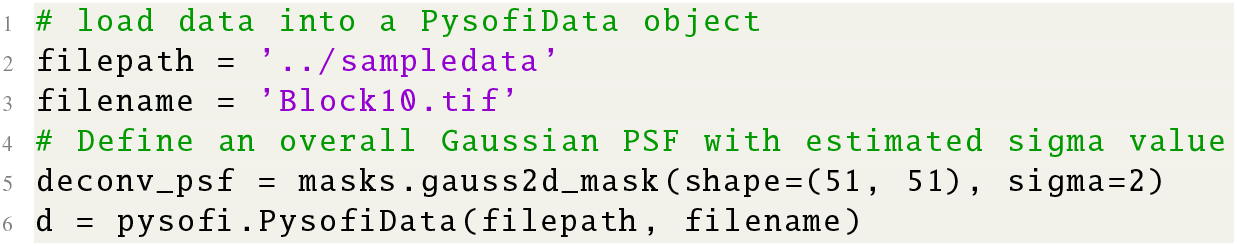
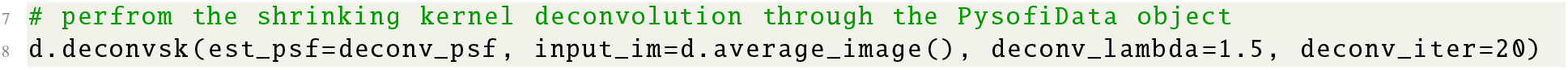

We can also use DeconvSK in *PySOFI* directly through the *deconvsk*.*py* module:

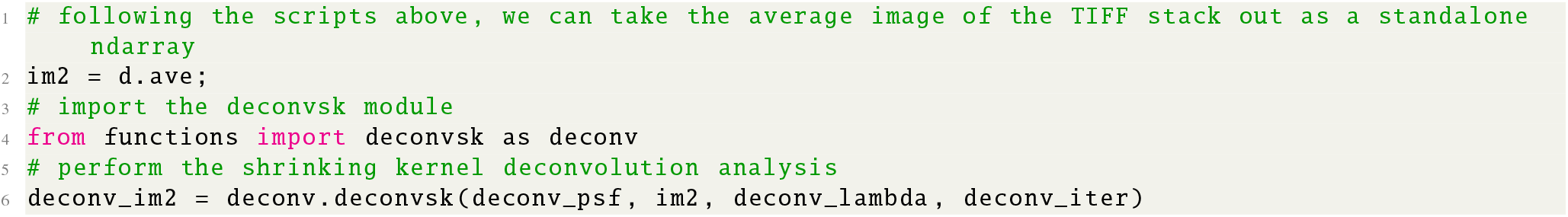

Figure 4 shows the deconvolution results on the relevant dataset used in our prior publication [11], we can see that the deconvolved results shows much more detailed structures as compared to the image without deconvolution. More detailed demonstrations are available in the corresponding Jupyter Notebook ([E3](https://github.com/xiyuyi-at-LLNL/pysofi/blob/master/Notebooks/E3_ShrinkingKernelDeconvolution.ipynb)).

**Figure 4:**
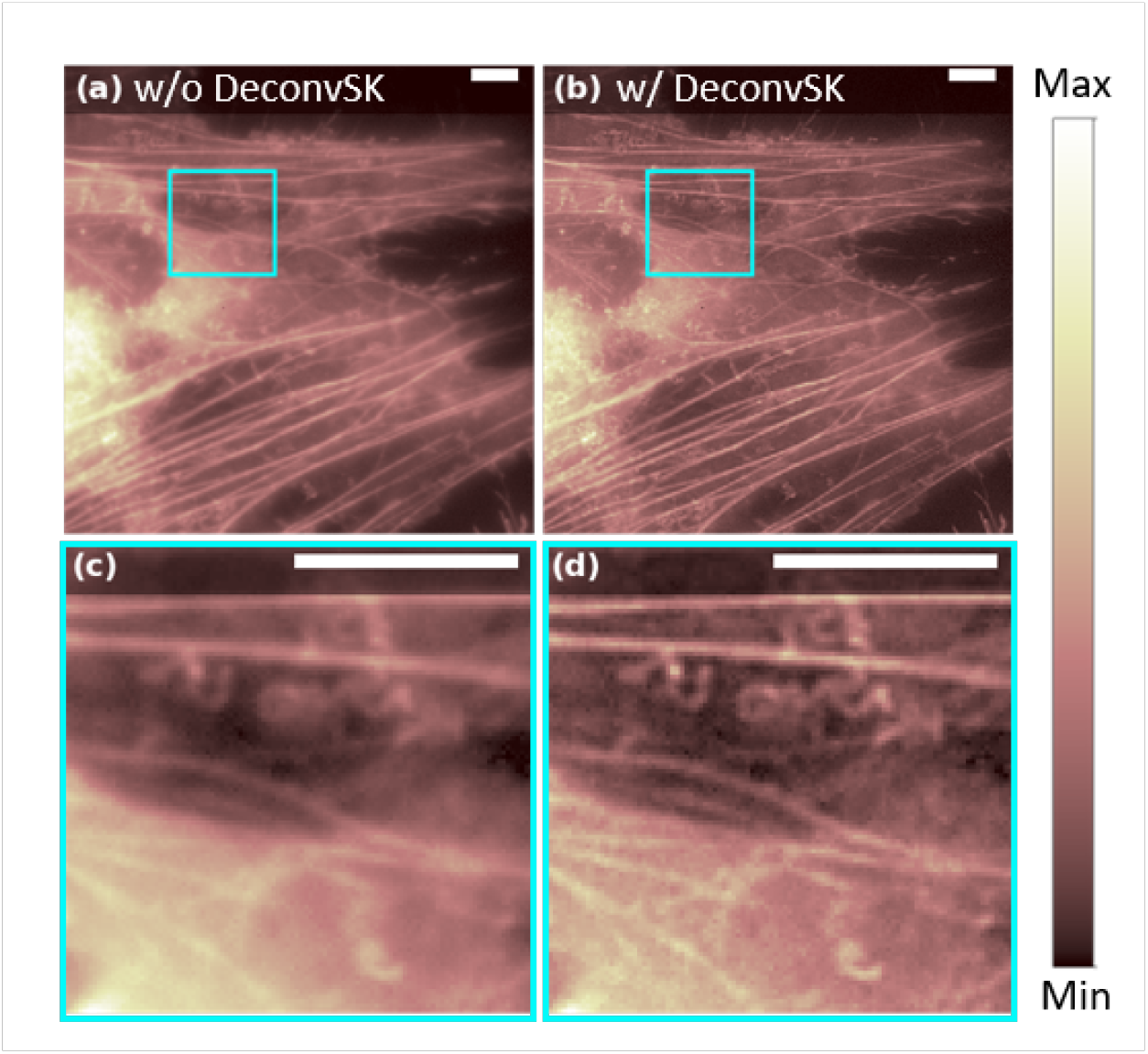
DeconvSK demonstration. Experimental demonstration of shrinking kernel deconvolution algorithm on HeLa cells transfected with Dronpa-C12 fused to *β*-Actin. Live cells were imaged with 30 ms frame integration and 200 frames in total. (a) Average widefield fluorescence image. (b) Average image after DeconvSK. (c) A zoom-in box of (a). (d) A zoom-in box of (b). Scale bars: 8*µm*.

### 3.4 Local dynamic range compression (*ldrc*) (E4)

One of the key challenges for high order SOFI cumulant calculations is the high dynamic range (HDR) of pixel intensities [1]. The HDR issue also exists in the high order moment images [11]. To mitigate such issues, We have developed the ‘local dynamic range compression (*ldrc*)’ method in our prior studies [11] and implemented it in the *PySOFI* package.

The *ldrc* algorithm rescales pixel intensities of a given image based on a reference image. First, a reference image with the same feature but a more confined dynamic range is defined (e.g., the average image, the second-order moment or cumulant SOFI image). The compression is performed locally in a small window that scans across the image with a stride of 1 pixel. In each window, the pixel intensities of the original image are linearly re-scaled to share the same dynamic range as the reference window [11]. The final value of each pixel is the average of the corresponding re-scaled values of them across all windows covering it.

In *PySOFI, ldrc* is implemented in the function module ldrc.py and integrated in the `PysofiData.ldrc()` method. The following scripts will calculate the 6^*th*^ order moment (*m6*) and the average image (*mean*), and perform *ldrc* on *m6* using *mean* as the reference:

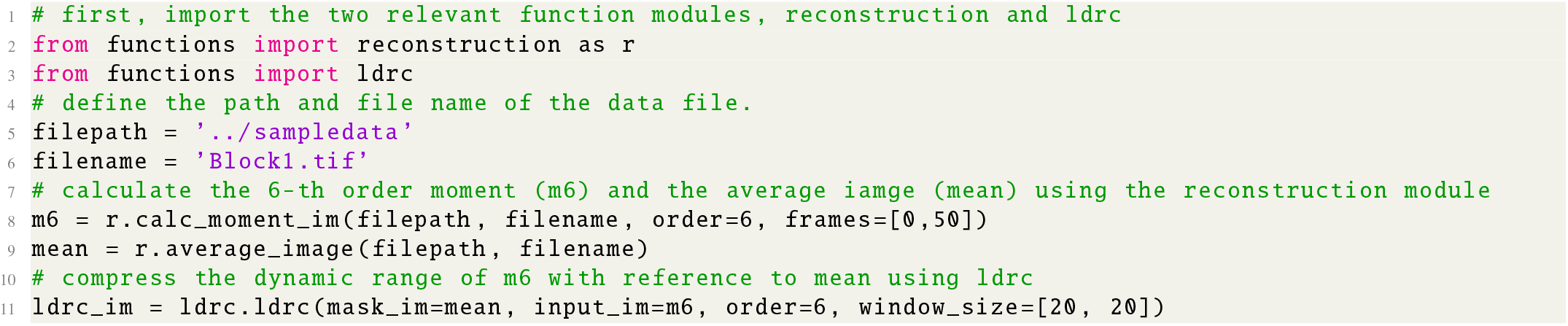

We can also perform the *ldrc* processing directly through the `PysofiData.ldrc()` method using the following script:

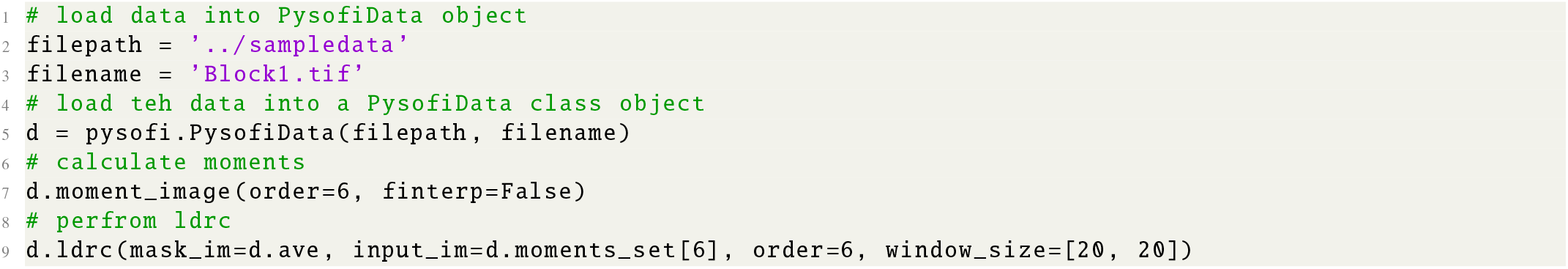

Note that the direct *ldrc* processing on *m6* often yields noisy results (We have demonstrated the results in the relevant Jupyter Notebook ([E4](https://github.com/xiyuyi-at-LLNL/pysofi/blob/master/Notebooks/E4_LDRCMethod.ipynb))). However, *ldrc* plays an important role in the SOFI 2.0 pipeline where the noise filtering and deconvolution are performed. In Figure 5, we compare the the partially processed SOFI 2.0 image (excluded *ldrc*) and the full SOFI 2.0 processed image (included *ldrc*). We can see that the feature in the image are preserved without *ldrc*, but imperceptible due to the HDR issue. On the other hand, *ldrc* mitigates the HDR issue and provide an image where the dim features are shown more clearly. More detailed demonstrations are available in the corresponding Jupyter Notebook ([E4](https://github.com/xiyuyi-at-LLNL/pysofi/blob/master/Notebooks/E4_LDRCMethod.ipynb)).

**Figure 5:**
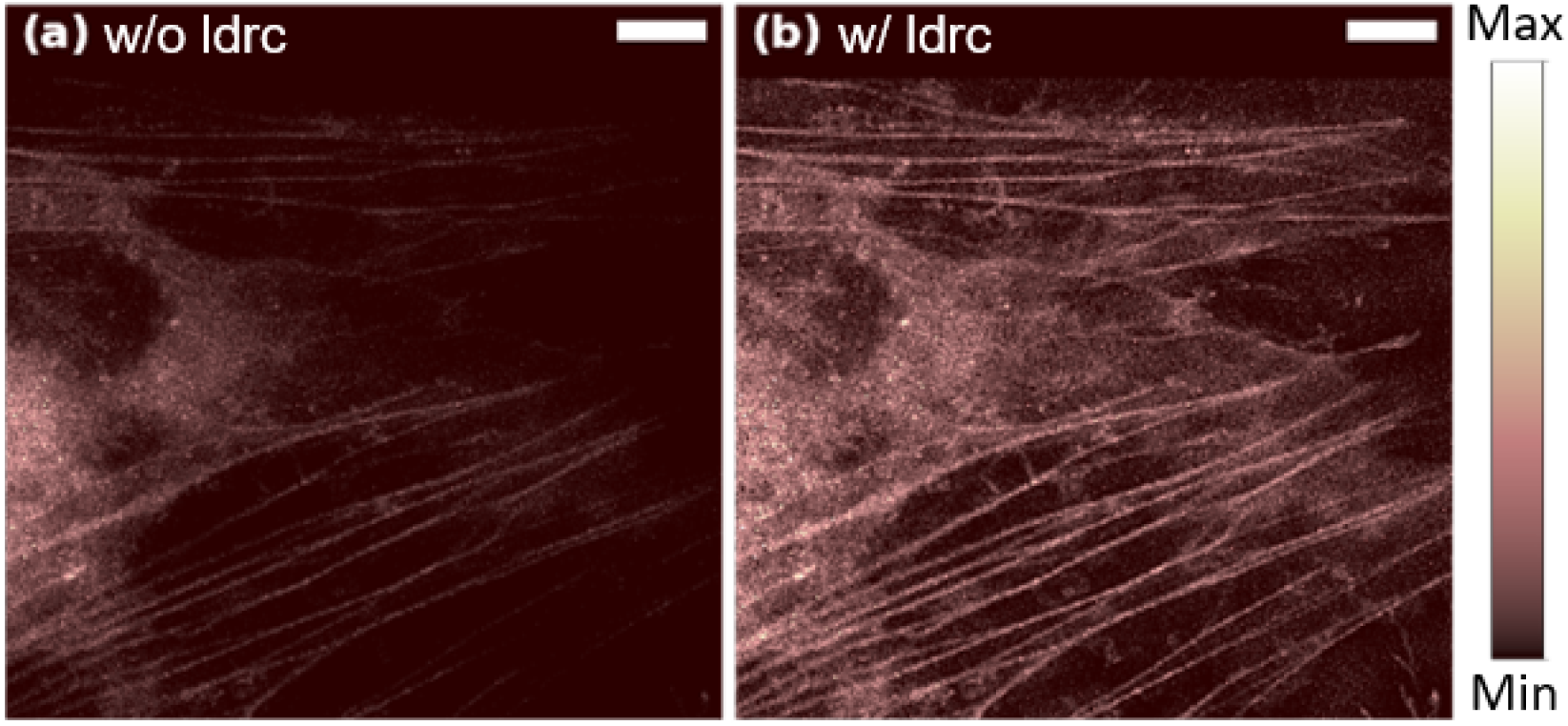
ldrc demonstration. Experimental demonstration of *ldrc* algorithm on HeLa cells transfected with Dronpa-C12 fused to *β*-Actin. Both images are processed using 6^*th*^ order moment, noise filtering and deconvolution, and obtained during the SOFI 2.0 analysis pipeline, before (a) and after (b) the *ldrc* step. Scale bars: 8*µm*.

### 3.5 Fourier interpolation (E5)

Fourier interpolation stochastic optical fluctuation imaging (fSOFI) solves the finite pixelation problem of SOFI by adding virtual pixels using Fourier transforms [9]. We have implemented the Fourier interpolation method in *PySOFI* to integrate the fSOFI analysis as an optional processing step. In our implementation, for the forward Fourier Transform, the Fourier transformation matrix was created with a size the same as the input image. We created the inverse Fourier transformation matrix to include the extra interpolation position coordinates, and omitted the “zero-padding” step in the Fourier space to avoid burdening the computation.With the Fourier interpolation, the input image/video is ‘projected’ onto a more refined grid with finer pixel size.

In *PySOFI*, Fourier interpolation is implemented in the function module finterp.py and integrated in the `PysofiData.finter p_tiffstack()` method. We can perform the Fourier interpolation and save the output as as a series of .tiff stacks. For example, the following scripts will calculate the 2- and 4-fold Fourier interpolation of the initial 100 frames from the example data set *block10*.*tiff*, and save the interpolated images into two .tiff stacks: `block10_InterpNum2.tiff` and `block10_InterpNum4.tiff` respectively.

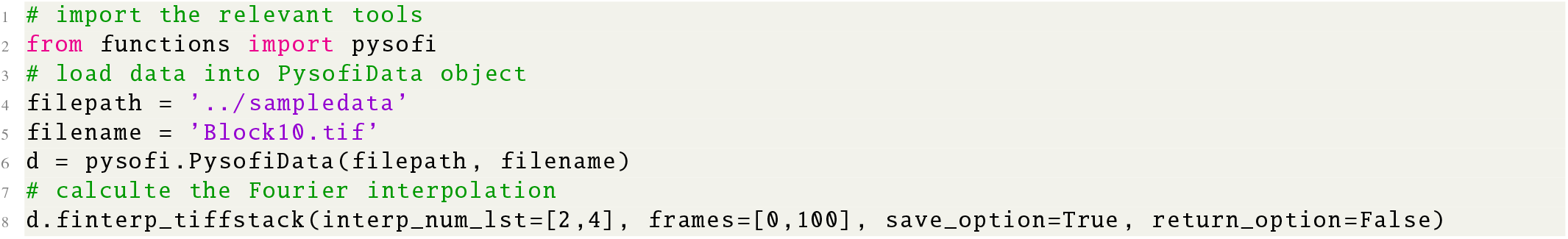

We can also perform the Fourier interpolation by using the `finterp.py` module as shown below:

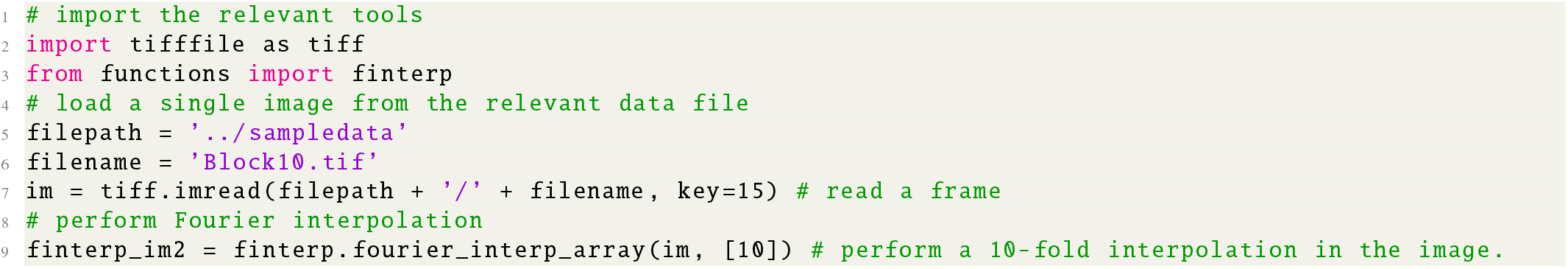

Figure 6 demonstrates the performance of the Fourier interpolation. Based on the Nyquist-Shannon sampling theorem [24, 25], we recommend setting the interpolation factor at least two times the highest order for moment-/cumulant reconstructions. For instance, if we plan to start the SOFI 2.0 pipeline with the 6^*th*^ order moment image, we should pass interp_num_lst = [12] to `d.finterp_tiffstack`. However, in practice, depending on the dimension and length of the input file, Fourier interpolation might consume large processing memory and time. If computation resources are limited, we recommend saving the interpolated image stack as tiff files firs tinstead of returning them, and then process the new file. Besides `d.finterp_tiffstack`, another option to include Fourier interpolation in the SOFI processing pipeline is to pass (`finterp = True`) and a interpolation factor (`interp_num=6`) when calculating the moment/cumulant reconstructions (see section 3.4).

**Figure 6:**
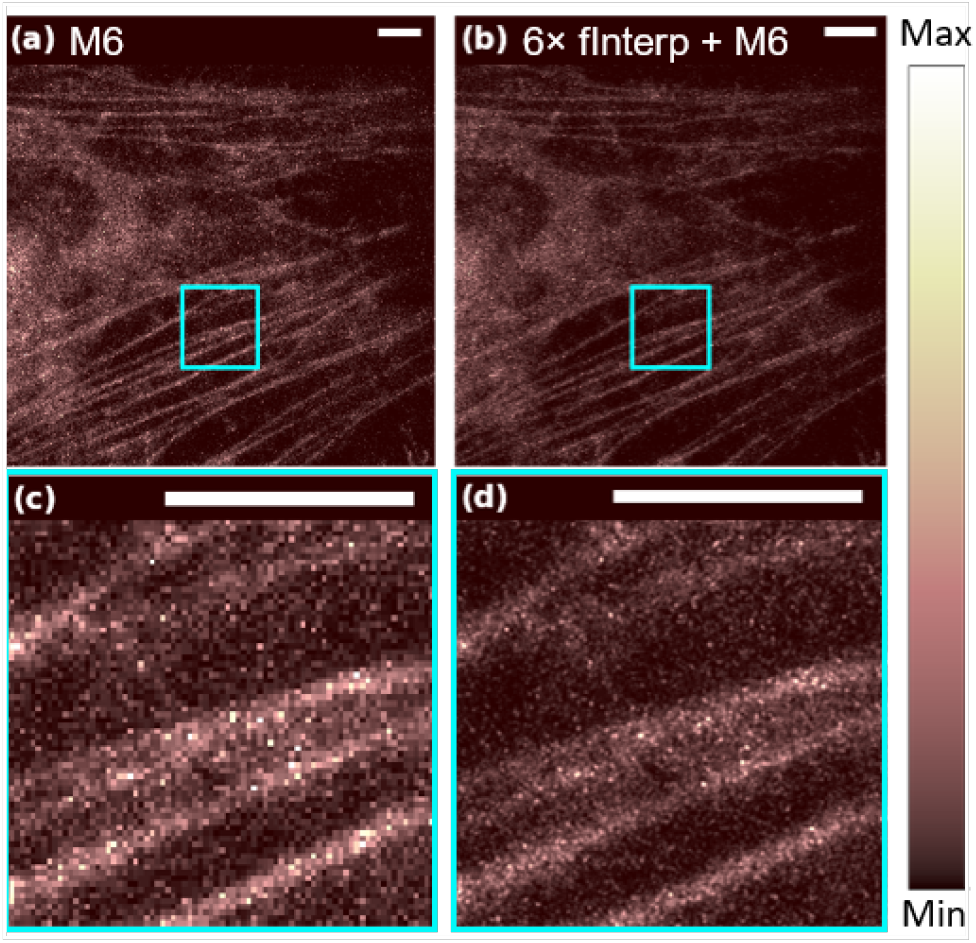
Fourier interpolation demonstration. Experimental demonstration of Fourier interpolation algorithm on HeLa cells transfected with Dronpa-C12 fused to *β*-Actin. (a) The 6^*th*^ order moment-reconstructed image of the original wide-field acquisition. (b) The 6^*th*^ order moment image after the Fourier interpolation. ldrc is performed on both (a) and (b) to compress the dynamic range of the reconstruction. (c) A zoom-in box of (a). (d) A zoom-in box of (d). Scale bars: 8*µm*.

More detailed demonstrations are available in the corresponding Jupyter Notebook ([E5](https://github.com/xiyuyi-at-LLNL/pysofi/blob/master/Notebooks/E5_FourierInterpolation.ipynb)).

### 3.6 Bleaching correction (E6)

Photobleaching of fluorescent probes is a general concern for super-resolution imaging analysis methods. As for SOFI, photobleaching can cause errors in virtual brightness displayed in moment or cumulant images [22]. Photobleaching leads to the loss of the fluorescence signal, which is mathematically equivalent as if the fluorophore is switched to a prolonged “off” state, degrading the quality of SOFI results. Therefore, a bleaching correction is critical.

*PySOFI* employs a bleaching correction technique [11] that divides the whole video into shorter blocks based on the total signal intensity, *I(t)*, where *t* is the time index, and *I(t)* is the summation of all the pixel values of the image at time index *t*. The individual blocks are processed independently and combined subsequently to form a SOFI movie. First, the time series of the total signal intensity is smoothened to obtain a monotonically decreasing curve as an estimation of the bleaching profile of the movie. Then, based on the signal evolution over time, the sizes of the shorter blocks are determined so that the fractional signal decrease within each block (characterized by the bleaching correction factor, *f*_*bc*_) is identical [11]. The final SOFI moment/cumulant images with bleaching correction are the average of those calculated from individual blocks. Fig. 7 shows that with the help of bleaching correction, the virtual brightness distribution and the photophysical properties (7c, 7f) are successfully restored, yielding similar values as compared to the simulated case without bleaching (7b).

**Figure 7:**
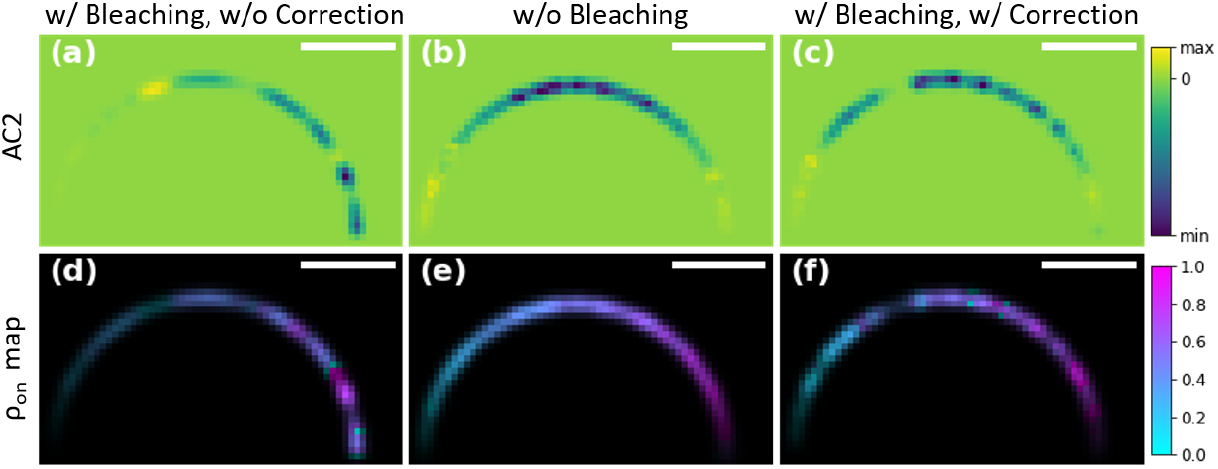
Bleaching correction demonstration on a simulation data. The fourth-order cumulant image (a-c) and multi-order cumulant analysis (MOCA) (d-f) is performed on a simulated video. A semicircle is populated with emitters with on-time ratios ranging from 0.01 (left) to 0.99 (right) with around 0.02 intervals. For emitters with photobleaching but without a bleaching correction step, the reconstructed pixel intensities (a) and emitters on-time ratio estimation (d) are far off from the true values (b, e), while the bleaching correction restores the information (c, f). Scale bars: 1.4*µm*.

*PySOFI* offers two ways for bleaching correction. One way is through the `PysofiData` class as shown below:

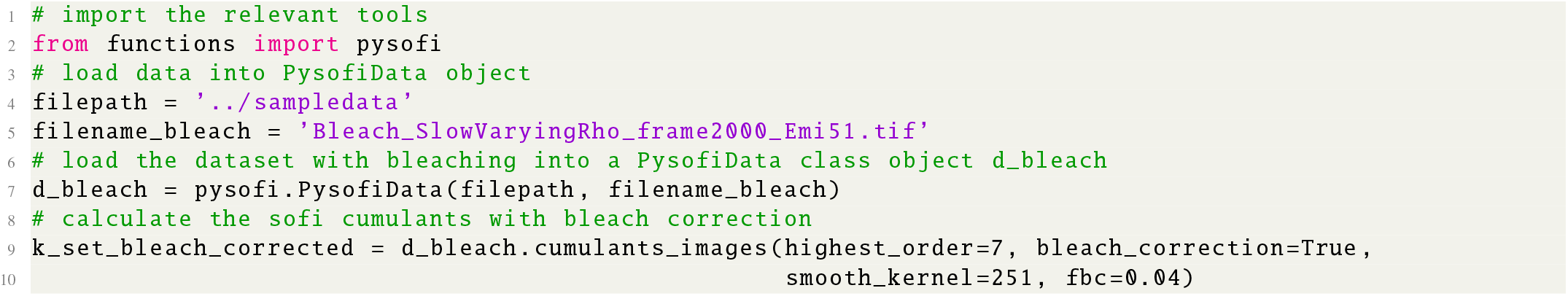

We can also directly import the relevant function module `reconstruction.py` and perfrom bleaching correction as shown below:

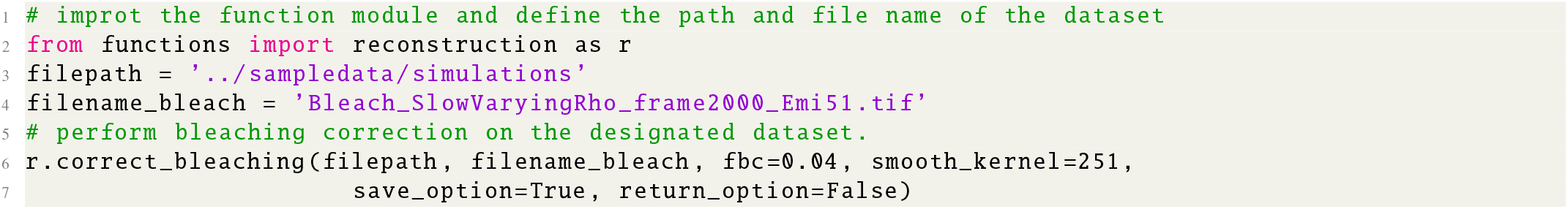

In this example, we applied bleaching correction to a TIFF stack, and the bleaching corrected movie is saved as a separate TIFF stack with the string “_bc” appended to the original file name.

More detailed demonstrations are available in the corresponding Jupyter Notebook ([E6](https://github.com/xiyuyi-at-LLNL/pysofi/blob/master/Notebooks/E6_BleachingCorrection.ipynb)).

### 3.7 Result visualization (E8)

We provide some simple visualization options in *PySOFI* to display either single or multiple images, with the options to adjust image contrast, and display the image with a transparency map defined as an input parameter. `Bokeh` is used to offer interactive display. More detailed demonstrations are available in the corresponding Jupyter Notebook ([E8](https://github.com/xiyuyi-at-LLNL/pysofi/blob/master/Notebooks/E8_ResultVisualization.ipynb)).

## 4. IMPLEMENTATION OF MULTI-ORDER CUMULANT ANALYSIS (MOCA)

This section describes multi-order cumulant analysis (MOCA) in *PySOFI*. The implementation is intended as an independent function module (*moca*.*py*) in the `functions` folder, providing an example for *PySOFI* extension.

MOCA characterizes and quantifies the photophysical properties of SOFI results [10]. It combines multiple orders of SOFI cumulants to construct a global fitting problem, and solves for the on-time ratio and on-state brightness for each pixel. A detailed description of MOCA can be found in [10]. In brief, MOCA is composed of two analysis steps. The first step is to estimate the PSF from the cross-correlation analysis. In the second step, MOCA reformulates multiple orders of cumulants and construct a global fitting problem to solve for the on-time ratio and on-state brightness. We briefly introduce the two analysis steps for MOCA below.

### 4.1. PSF estimation from SOFI auto-/cross-cumulants

The *n*^*th*^-order SOFI auto-cumulants 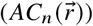 are images where the value of a pixel at location 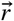 is the cumulant value calculated from a single-pixel corresponding to the same position in the input data. Cross-cumulants 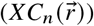 are images where a pixel at position 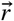 is calculated based on the cross-cumulants of a group of different pixels whose geometric center is located at the image pixel position 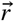. Intrinsically, the choice of the group of pixels used to calculate a SOFI-*XC*_*n*_ pixel can either be *n* distinct pixels or less than *n* pixels where a subset of the pixels can be used more than once [8]. By combining different orders of SOFI-*AC*_*n*_ and different orders of SOFI-*XC*_*n*_ with different choices of pixel combinations, one can achieve a robust estimation of the PSF through a global fitting.

We start with the mathematical expression of the *n*^*th*^*-*order SOFI auto-cumulants *AC*_*n*_ and SOFI cross-cumulants *XC*_*n*_ (with all the time lags equal to 0):

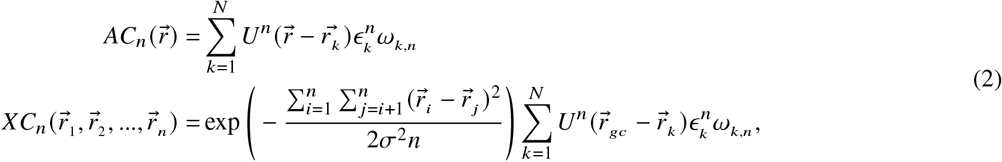

where *k* is the emitter index, *ϵ*_*k*_ is the brightness of on-state of emitter *k* and *ω*_*k*_ is the *n*^*th*^ order cumulant of the blinking profile of the *k*^*th*^ emitter. *U* is the PSF of the optical system that can be assumed as a 2D Gaussian function 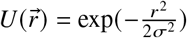 (we consider 2D images here). 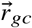 is the geometric center of set of pixels used for the cross-cumulant calculations, for which we have 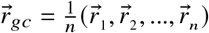. If we carefully choose the set of pixels to have geometric center located at 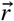, we would have the following:

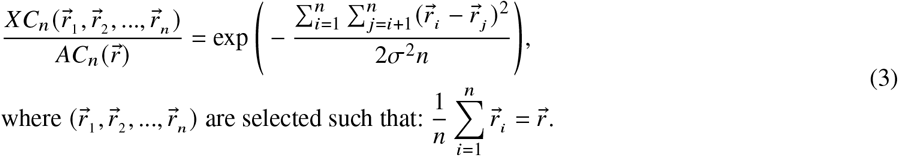

We define 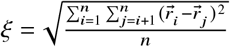 to simplify the notation into the following:

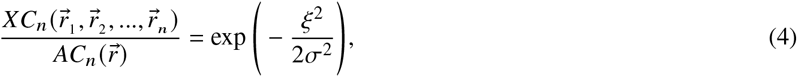

where *XC*_*n*_ and *AC*_*n*_ are computed from the imaging data, and *ξ* can be calculated based on the choices of the pixel combinations. The only unknown is *σ* that characterizes the size of PSF. In this case, multiple pixel combinations can be selected to create different 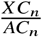 and *ξ* ^2^ values, and a global fitting can then be applied to estimate the PSF (*σ*). We demonstrate with *n*= 2 in this manuscript, but the principle applies to any order and can be implemented when necessary. Details of the global fitting approach are explained in chapter 7 of [10].

### 4.2. Local parameter mapping from MOCA

Assume that the spatial variance of the photophysical properties of the fluorophores are small enough to be invariant within the range of the spatial resolution of the lowest order cumulant used [22],*n*^*th*^ order SOFI auto-cumulants can be simplified by removing the emitter indices and expressing the relevant physical quantities as spatial variables:

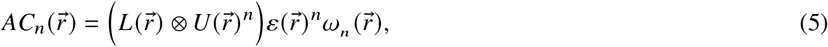

where 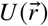 is the PSF, 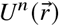 is the virtual PSF for *n*^*th*^ order SOFI images, and ⊗ indicates spatial convolution operator. 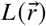 encodes the location information of emitters, for which we have 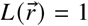 if there is an emitter at location 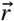, and 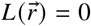 otherwise. *ϵ*_*k*_ and *ω*_*k,n*_ in equation 2 are assumed to have small spatial variations where emitter brightness and blinking statistics are primarily dependent on the emitters’ local environment (such as PH level or oxygen level). Thus, in equation 5 we replace the notation of *ϵ*_*k*_ into 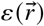 that describes the summation of the on-time brightness of emitters at 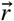. Similarly, *ω*_*k,n*_ is replaced as 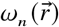 to describe the blinking statistics of emitters at 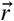. Note that *ω*_*k,n*_ is a polynomial function of the on-time ratio *ρ*_*on*_ (shown in equation 4.2 in [22]). With the reformatted equation 5, 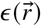 and 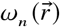 encode the physical properties of the emitters as an environment-dependent spatial variable.

We can then define a new quantity, *X*_*n*_, to connect multiple orders of SOFI auto-cumulants by convolving both sides of equation 5 with 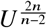 followed by dividing with *AC*_2_:

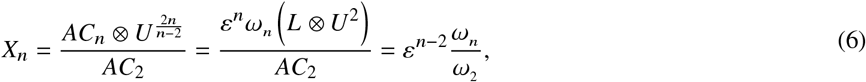

Specifically, the numerical values of *X*_*n*_ can be calculated from different orders of auto-cumulants. Meanwhile, as shown in equation 6, we know that *X*_*n*_ is a function of *ε* and *ρ*_*on*_ (because *ω*_*n*_ is a function of *ρ*_*on*_). Therefore, we can calculate various orders of *X*_*n*_ to obtain an equation system for which the only unknown variables are *ε* and *ρ*_*on*_, and any regression method that solves a polynomial equation system with 2 unknown variables can solve for the spatial variables *ε* and *ρ*_*on*_

Here we solve the equation system by reformatting equation 6 further:

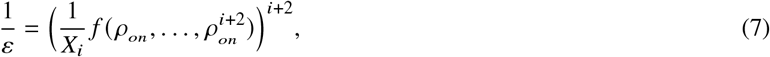

which describes a curve in a 2D coordinate system with axes 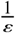 and *ρ*_*on*_ .We calculate auto-cumulants up to the 7^*th*^order, and use *X*_3_ ∼ *X*_7_ (five equations, five curves) to find the global optimal solution for 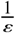 and *ρ*_*on*._ Note that *ρ*_*on*_ (x-axis) is naturally bounded between 0 to 1, and *ε* is naturally positive so 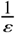 (y-axis) is bounded between 0 and 1, therefore we have a bounded 2D space to search for the optimum solution.

Ideally, the common crossing point of multiple curves presented by the equation system would be the solution. But due to numerical errors and imperfections of the model, the five curves wouldn’t have a common crossing point. In our implementation, we first define a quantity *D*(*ρ*_*on*_) as the sum of mutual distances square of 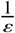 (y-axis) among the points on all the curves at each *ρ*_*on*_ value, which can characterize the dispersion of those points. We sweep through the *ρ*_*on*_ values (x-axis) to calculate *D*(*ρ*_*on*_), and determine the final solution of *ρ*_*on*_ to be the one that yields the smallest *D*(*ρ*_*on*_) value. The final solution of 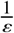 is determined by the average of the y-coordinates among all the identified crossing points. We fit such equation system for every pixel to map 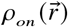 and 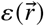 values to obtain the spatial distribution of the photo-physical properties of the emitters.

### 4.3. Use MOCA with *PySOFI* (E7)

MOCA is implemented as an extension module supported by existing *PySOFI* modules and functions. Here, MOCA methods are imported as the `moca.py` module directly as shown below:

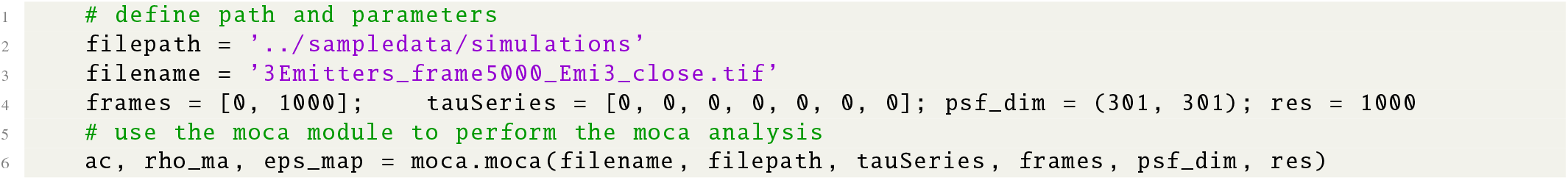

To demonstrate the performance of MOCA under a complex condition, we created a simulated data set with 18 curves randomly decorated with blinking emitters. The *ρ* value of each curve is selected among 0,3, 0.5 and 0.7 (six curves for each *ρ* value). Among six curves with the same *ρ* value, two of them are 2 times brighter than the other four. As shown in figure 8, MOCA successfully recovered different on-time ratio and emitter brightness under various conditions (e.g., intersections of filaments with emitters with different photophysical properties.

**Figure 8:**
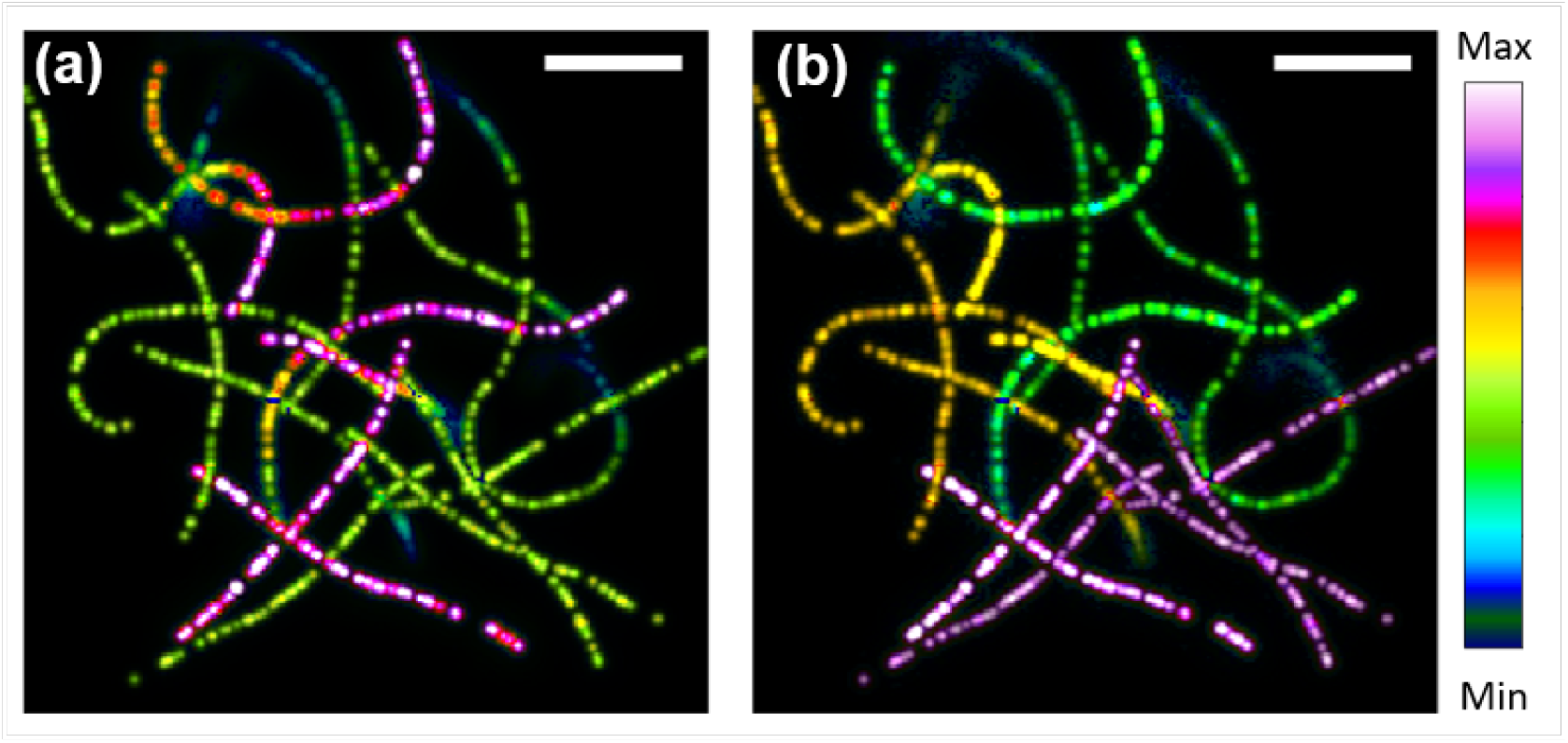
MOCA demonstration on simulated filaments. Simulated crossing filaments with different *ρ*’s are generated. There are three *ρ*’s, 0.3, 0.5 and 0.7, and for each *ρ*, there are six curves. MOCA estimates the on-time ratio and brightness of emitters precisely even at the intersections. Scale bar: 7*µm*.

More examples are available in the corresponding Jupyter Notebook ([E7](https://github.com/xiyuyi-at-LLNL/pysofi/blob/master/Notebooks/E7_MOCA.ipynb)).

## 5. DISCUSSION

In this work, we developed *PySOFI*, an open source python package for SOFI analyses. *PySOFI* contains the essential functionalities for conventional SOFI analysis as well as several derivative methods [9, 11, 22, 10].

PySOFI adopts a simple architecture, where all the data processing steps are implemented as independent function modules, and only one class module (the data class `PysofiData`) is used to manage the data processing workflow. The functions can be tested independently and used in different processing pipelines. A fast prototype on new analysis can be achieved by disseminating and reorganizing the processing step. Like the example of MOCA as a *PySOFI* extension module in section 4, one can implement additional processing steps as independent python functions with the help of existing *PySOFI* functions. New functions can be used as standalone modules, or can be integrated into the `PysofiData` class to support the new analysis pipeline. New classes can be constructed for different analysis pipelines as well.

We adopted [Sphinx](https://www.sphinx-doc.org/en/master/) to manage the *PySOFI* documentation, which is available as an [online documentation](https://xiyuyi-at-llnl.github.io/pysofi/build/html/) to facilitate community usage. Additionally, each processing element of the analysing pipeline are demonstrated in individual [Jupyter Notebooks](https://github.com/xiyuyi-at-LLNL/pysofi/tree/master/Notebooks). In each notebook, we also provide instructions on how to tune processing variables and explore input data.

*PySOFI* is housed on [GitHub](https://github.com/xiyuyi-at-LLNL/pysofi) as an open source repository, any interested individuals can learn, inspect, validate and contribute to the package. The user interactions on GitHub (e.g., fork, create pull requests, and report issues) engage the community communications. We expect *PySOFI* to benefit general SOFI users for existing SOFI analysis, as well as developers and new investigators interested in developing new SOFI-relevant analysis method.

## ACKNOWLEDGMENTS

The work from Y. M. and S. W. was supported by the National Science Foundation under Grant No. DMR-1548924 and by the Dean Willard Chair funds. The work from X. Y. was performed under the auspices of the U.S. Department of Energy by Lawrence Livermore National Laboratory under Contract DE-AC52-07NA27344. Release number: LLNL-JRNL-827376.

## AUTHOR CONTRIBUTIONS

Y.M. designed the *pysofi* architecture, implemented the *pysofi* package, generated simulation videos for figure 7. X.Y designed the research, supervised the development of *PySOFI* package, developed the MOCA algorithms and collected experimental and simulation data. All authors analyzed the data, discussed the results, and wrote the manuscript.

## DATA AVAILABILITY

The data for this project is partially available on the [project repository](https://github.com/xiyuyi-at-LLNL/pysofi), and partially available on [figshare](https://figshare.com/s/47d97a2df930380c96bb).

## REFERENCES

[1] Thomas Dertinger et al. “Fast, background-free, 3D super-resolution optical fluctuation imaging (SOFI)”. In: Proceedings of the National Academy of Sciences 106.52 (2009), pp. 22287–22292.

[2] Kristin Grußmayer et al. “Self-Blinking Dyes Unlock High-Order and Multiplane Super-Resolution Optical Fluctuation Imaging”. In: ACS nano 14.7 (2020), pp. 9156–9165.

[3] S Duwé, W Vandenberg, and P Dedecker. “Live-cell monochromatic dual-label sub-diffraction microscopy by mt-pcSOFI”. In: Chemical Communications 53.53 (2017), pp. 7242–7245.

[4] Benjamien Moeyaert and Peter Dedecker. “PcSOFI as a smart label-based superresolution microscopy technique”. In: Photoswitching Proteins. Springer, 2014, pp. 261–276.

[5] Sam Duwé, Benjamien Moeyaert, and Peter Dedecker. “Diffraction-Unlimited Fluorescence Microscopy of Living Biological Samples Using pcSOFI”. In: Current protocols in chemical biology 7.1 (2015), pp. 27–41.

[6] Peter Dedecker, Gary CH Mo, and Jin Zhang. “Widely Accessible Method for Superresolution Fluorescence Imaging of Living Systems”. In: Biophysical Journal 104.2 (2013), 535a.

[7] Stefan Geissbuehler et al. “Mapping molecular statistics with balanced super-resolution optical fluctuation imaging (bSOFI)”. In: Optical Nanoscopy 1.1 (2012), pp. 1–7.

[8] Thomas Dertinger et al. “Achieving increased resolution and more pixels with Superresolution Optical Fluctuation Imaging (SOFI)”. In: Optics express 18.18 (2010), pp. 18875–18885.

[9] Simon C Stein et al. “Fourier interpolation stochastic optical fluctuation imaging”. In: Optics express 23.12 (2015), pp. 16154–16163.

[10] Xiyu Yi. “Super resolution of Optical Fluctuation Imaging 2.0 (SOFI-2.0): Towards fast super resolved imaging of live cells”. PhD thesis. UCLA, 2017.

[11] Xiyu Yi et al. “Moments reconstruction and local dynamic range compression of high order Superresolution Optical Fluctuation Imaging”. In: Biomedical optics express 10.5 (2019), pp. 2430–2445.

[12] S Vlasenko et al. “Optimal correlation order in superresolution optical fluctuation microscopy”. In: Physical Review A 102.6 (2020), p. 063507.

[13] Dario Cevoli et al. “Design of experiments for the optimization of SOFI super-resolution microscopy imaging”. In: Biomedical Optics Express 12.5 (2021), pp. 2617–2630.

[14] Baoju Wang et al. “Active-modulated, random-illumination, super-resolution optical fluctuation imaging”. In: Nanoscale 12.32 (2020), pp. 16864–16874.

[15] Adrien Descloux et al. “Combined multi-plane phase retrieval and super-resolution optical fluctuation imaging for 4D cell microscopy”. In: Nature Photonics 12.3 (2018), pp. 165–172.

[16] Eliel Hojman et al. “Photoacoustic imaging beyond the acoustic diffraction-limit with dynamic speckle illumination and sparse joint support recovery”. In: Optics express 25.5 (2017), pp. 4875–4886.

[17] MinKwan Kim et al. “Superresolution imaging with optical fluctuation using speckle patterns illumination”. In: Scientific reports 5.1 (2015), pp. 1–10.

[18] Ori Katz and Noam Shekel. “Using fiber-bending generated speckles for improved working distance and”. In: Optics Letters 4.C1 (2020), p. C2.

[19] Lydia Kisley et al. “Characterization of porous materials by fluorescence correlation spectroscopy super-resolution optical fluctuation imaging”. In: ACS nano 9.9 (2015), pp. 9158–9166.

[20] Stefan Geissbuehler et al. “Live-cell multiplane three-dimensional super-resolution optical fluctuation imaging”. In: Nature communications 5.1 (2014), pp. 1–7.

[21] Ashley M Rozario et al. “Corrigendum to:’Live and Large’: Super-Resolution Optical Fluctuation Imaging (SOFI) and Expansion Microscopy (ExM) of Microtubule Remodelling by Rabies Virus P Protein”. In: Australian Journal of Chemistry 73.8 (2020), pp. 822–822.

[22] Xiyu Yi and Shimon Weiss. “Cusp-artifacts in high order superresolution optical fluctuation imaging”. In: Biomedical optics express 11.2 (2020), pp. 554–570.

[23] Peter Dedecker et al. “Localizer: fast, accurate, open-source, and modular software package for superresolution microscopy”. In: Journal of biomedical optics 17.12 (2012), p. 126008.

[24] Harry Nyquist. “Certain topics in telegraph transmission theory”. In: Transactions of the American Institute of Electrical Engineers 47.2 (1928), pp. 617–644.

[25] Claude E Shannon. “A mathematical theory of communication”. In: The Bell system technical journal 27.3 (1948), pp. 379–423.

